# Nuclear activities and interactome of the NS5 protein of Tick-Borne Encephalitis Virus

**DOI:** 10.1101/2025.11.29.691275

**Authors:** Maxime Chazal, Aïssatou Aïcha Sow, Elodie Le Seac’h, Dana Hawasheen, Margarida Bonifacio, Mikael Feracci, Ségolène Gracias, Adrià Sogues, Melissa Molho, Heng Pan, Boris Bonaventure, Jean-Pierre Quivy, Geneviève Almouzni, Etienne Decroly, Holly Ramage, Jeffrey R. Johnson, Vincent Caval, Nolwenn Jouvenet

**Affiliations:** Unité Signalisation Antivirale, UMR CNRS 3569, Université Paris Cité, Institut Pasteur, Paris, France; Aix Marseille Université, AFMB, UMR CNRS 7257, Marseille, France; Bacterial Cell Cycle Mechanisms Unit, UMR CNRS 3528, Université Paris Cité, Institut Pasteur Paris, France; Department of Microbiology and Immunology, Thomas Jefferson University, Philadelphia, USA; Department of Microbiology, Icahn School of Medicine at Mount Sinai, New York, USA; Global Health and Emerging Pathogens Institute, Icahn School of Medicine at Mount Sinai, New York, USA; Institut Curie, PSL Research University, Sorbonne Université, UMR CNRS 3664, Laboratoire Dynamique du Noyau, Equipe Labellisée Ligue contre le Cancer, Paris, France

## Abstract

Orthoflaviviruses are RNA viruses responsible for significant diseases in humans, domesticated animals and wildlife. Their NS5 protein is central in viral replication, functioning both as an RNA-dependent RNA polymerase and a methyltransferase, while also modulating cellular processes, including the interferon response. Although viral replication is cytoplasmic, the NS5 protein of several mosquito-borne orthoflaviviruses cycles between the cytoplasm and the nucleus of infected human cells. However, the nuclear localization and function of NS5 of tick-borne orthoflaviviruses, such as tick-borne encephalitis virus (TBEV), remained poorly understood. Microscopy analysis and cell fractionation revealed that the NS5 protein of TBEV localized to both the cytoplasm and nucleoplasm of infected cells. Mutagenesis studies identified critical residues required for its nuclear targeting. Mutating these residues in a TBEV replicon abolished viral replication. Immunoprecipitation-mass spectrometry analyses performed in two human cell lines infected with TBEV recovered 352 NS5 partners. Among them, 187 were nuclear or partially nuclear. By integrating our interactome data with that of Powassan virus (POWV), another tick-borne orthoflavivirus, we refined a list of 20 high-confidence NS5 partners, including splicing factors and chromatin modulators. Functional analysis revealed that seven of these nuclear partners significantly modulated viral replication, further underscoring the importance of nuclear NS5 in the viral life cycle. Our work advances our understanding of the nuclear function of the NS5 proteins of tick-borne orthoflaviviruses.

**Importance:** Tick-borne orthoflaviviruses are emerging globally, spreading across Europe, Asia, and North America, where they infect humans, domesticated animals, and wildlife. These viruses produce a protein called NS5, which drives viral replication and helps evade the innate immune response. We observed that the NS5 protein of tick-borne encephalitis virus (TBEV) localized both in the cytoplasm and nucleoplasm of infected human cells. We identified the specific residues responsible for its nuclear addressing and showed that it interacts with numerous nuclear proteins, including some involved in regulating gene expression. Seven of these nuclear partners significantly influenced viral replication, highlighting the importance of NS5’s nuclear activity. This work sheds light on how tick-borne orthoflaviviruses manipulate host cells, deepening our understanding of their replication strategies.

## Introduction

Most orthoflaviviruses are primarily transmitted to vertebrate hosts by mosquitoes or ticks [1]. They have a world-wide distribution and infect up to 400 million people annually. They cause a wide spectrum of diseases, including hepatitis, vascular leakage, shock syndrome, encephalitis, acute flaccid paralysis, congenital abnormalities, and death [1]. Over the past 50 years, outbreaks of orthoflaviviruses have increased significantly. For instance, the introduction into the Western Hemisphere of West Nile virus (WNV) in 1999 and Zika virus (ZIKV) in 2015 have led to large numbers of severe human infections [2,3]. Dengue fever, which is caused by one of the four serotypes of dengue virus (DENV), is currently experiencing an unprecedented worldwide resurgence [4]. Meanwhile, Yellow fever virus (YFV) outbreaks, driven by insufficient vaccination coverage, remain a major concern [5]. Finally, tick-borne orthoflaviviruses are also emerging in various parts of the world, as seen with tick-borne encephalitis virus (TBEV) in Europe and Powassan virus (POWV) in North America [6].

Orthoflaviviruses share a common genome organization and coding strategy [1]. Upon viral entry into the host cell and fusion between the viral and endosomal membranes, the 11-kb viral genome is released into the cytosol and addressed to the endoplasmic reticulum (ER) to be translated by the cellular machinery into a large polyprotein precursor. This polyprotein is processed by host and viral proteases into three structural proteins (the capsid protein (C), the precursor of the M protein (prM) and the envelope (E) glycoprotein) and seven non-structural (NS) proteins (NS1, NS2A, NS2B, NS3, NS4A, NS4B and NS5) [1]. The structural proteins form the virus particles whereas the NS proteins are involved in viral replication and assembly, as well as in counteracting the cellular antiviral response [1].

The orthoflavivirus NS5 protein is a highly conserved protein of about 100 kDa. It contains two enzymatic domains, an N-terminal methyltransferase (MTase) domain and an RNA-dependent RNA polymerase (RdRp) domain, which are responsible for capping and replication of the viral genome, respectively [7]. NS5 is also a potent inhibitor of interferon (IFN) signaling [8,9]. Although viral replication occurs within the cytoplasm, the NS5 protein of several mosquito-borne orthoflaviviruses, including DENV and ZIKV, localizes to the nucleus of infected human cells [10–12]. DENV NS5 carries two highly conserved nuclear localization signals (NLS) that are recognized by the importin β1 and importin α/β nuclear import factors [10,13], with the αβNLS likely being the primary determinant of nuclear addressing [10]. ZIKV NS5, which localizes mainly in punctate nuclear bodies in infected cells, also interacts with importin β1 and importin α/β [11,12].

Several functions have been attributed to the nuclear localization of the NS5 protein of mosquito-borne orthoflaviviruses. Nuclear translocation may protect NS5 from proteolytic degradation in the cytoplasm [11]. Nuclear NS5 may also exert proviral functions *via* interactions with nuclear partners. Indeed, proteomic analysis coupled to functional studies in human hepatocarcinoma Huh7 cells and embryonic kidney 293T cells infected with DENV2 revealed that NS5 perturbs the correct splicing of antiviral factors by interacting with core components of the splicing machinery [14]. Similarly, proteomic analysis using 293T cells expressing tagged versions of the NS5 protein of ZIKV, DENV2, WNV and Japanese encephalitis virus (JEV) recovered numerous spliceosomal components and spliceosome-associated proteins [15–17], suggesting that NS5-mediated targeting of the spliceosome is a strategy shared by mosquito-borne orthoflaviviruses to modulate gene expression. Finally, ZIKV NS5 blocks the transcriptional elongation of numerous genes *via* a direct binding to chromatin, leading to a disruption of neurogenesis in mice brain [18]. In tick-borne orthoflaviviruses, the nuclear localization of NS5 and its functional consequences remain unclear.

We recently produced an antibody against TBEV NS5 and observed, using confocal microscopy analysis, that the protein localized both in the cytoplasm and nucleoplasm of infected Huh7 cells [9]. We aimed here to better understand what controls the nuclear presence of TBEV NS5 and get insight into its functional role. Using mutagenesis approaches and immunoprecipitation-mass spectrometry (IP-MS) of infected cells coupled to functional studies, we identified residues critical for NS5 nuclear localization and provided a list of NS5 nuclear partners, some of which modulate viral replication.

## Results

### The nuclear accumulation of TBEV NS5 requires importin β1- and α/β-interacting domains

We previously observed, using an in-house generated TBEV NS5 antibody, that TBEV NS5 localized both in the cytoplasm and nucleus of Huh7 cells infected for 48 hours with TBEV (Hypr strain) at an MOI of 0.02 [9]. We confirmed the localization of NS5 using human microglial cells (HMC3), which are relevant for TBEV biology since they are sites of viral replication *in vivo* [19]. As in Huh7 cells [9], TBEV NS5 exhibited a diffuse cytoplasmic and nuclear localization in HMC3 cells infected at an MOI of 1 for 24 hours (Fig. 1A and S1A). The NS5 signal was excluded from the nucleolus in infected HCM3 (Fig. 1A) and Huh7 cells (Fig. S1B). By contrast, the E signal was exclusively cytoplasmic (Fig. 1A).

**Figure 1.**
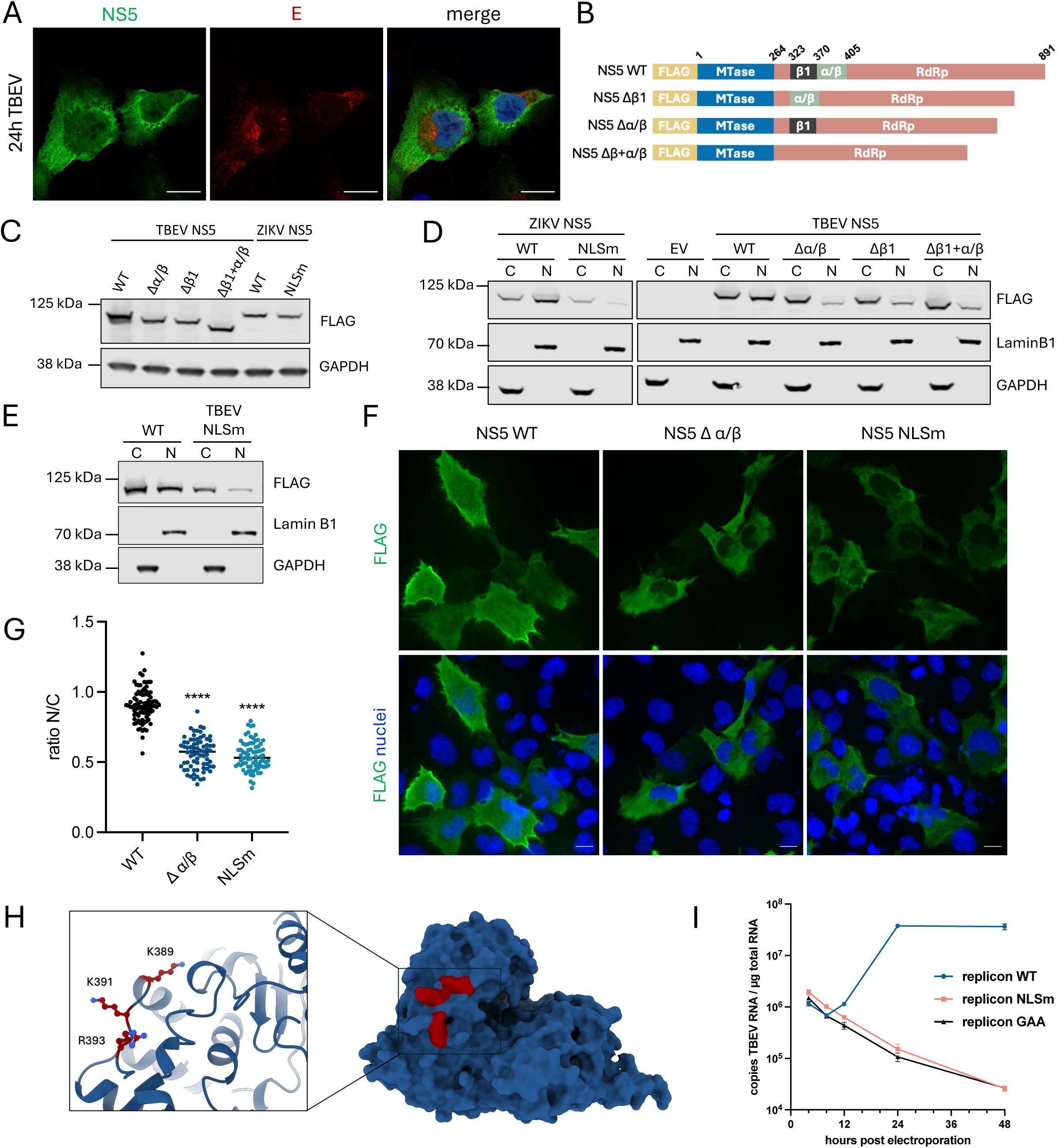
Characterization of TBEV NS5 nuclear localization. A. HMC3 cells were infected with TBEV at an MOI of 1. After 24 hours, subcellular localization of NS5 was determined by immunofluorescence microscopy using antibodies against NS5 (green) or E (red). Nuclei were stained using DAPI (blue) and images were acquired using a confocal microscope. Scale bars 20 µm. B. Schematic representation of wild-type (WT) and mutants NS5 proteins. C. 293T cells were transfected with plasmids encoding TBEV or ZIKV FLAG-NS5 WT or mutants. After 24 hours, whole-cell lysates were analyzed by western blot using antibodies against FLAG and GAPDH. D. E. 293T cells were transfected with plasmids encoding TBEV or ZIKV FLAG-NS5, WT or mutants. After 24 hours, NS5 subcellular localization was determined by cellular fractionation. Cytoplasmic (C) and nuclear (N) fractions were analyzed by western blot using antibodies against FLAG, GAPDH and Lamin B1. F. HMC3 cells were transfected with plasmids expressing FLAG-tagged NS5 WT, Δα/β or NLSm. After 24 hours, subcellular localization of NS5 was determined by immunofluorescence microscopy using an antibody recognizing FLAG (green). Nuclei were stained using DAPI (blue) and images were acquired using a brightfield microscope. Scale bars 10 µm. G. Quantification of the nuclear localization of NS5 WT or mutants was determined by measuring the intensity of cytoplasmic (C) and nuclear (N) FLAG signal from images shown in F. using Fiji. The ratio of mean fluorescence N/C is represented. 73 to 85 cells were analyzed for each condition, ****p < 0.0001, unpaired t-test. H. AlphaFold model of the RdRp domain of TBEV NS5. The three amino acids essential for nuclear import are highlighted in red. I. The effect of the NS5 nuclear localization on viral replication was assessed by mutating the NLS signal in a TBEV replicon. *In vitro* transcribed RNAs from WT or mutated replicons were electroporated into HMC3 cells and viral replication was measured using RT-qPCR.

The nuclear localization signals (NLS) of the NS5 proteins of several mosquito-borne orthoflaviviruses have been characterized. Both DENV and ZIKV NS5 carry a bipartite NLS that interacts with importin α/β and importin β1, with the importin α/βNLS being the predominant one for DENV2 NS5 [10–13]. Using alignment of the NS5 proteins of DENV2, WNV, YFV, ZIKV and TBEV (Fig. S1C), we found that TBEV NS5 also contains basic residues within the N-terminal region of its RdRp domain that may bind importin β1 and importin α/β and thus could serve as an NLS. To determine the impact of these 2 importin-binding domains in TBEV NS5 nuclear accumulation, we generated versions of the protein lacking either importin β1-binding residues (Δβ1), importin α/β-binding residues (Δα/β), or both domains (Δβ1+α/β). Wild-type TBEV NS5 and the three deletion mutants were FLAG-tagged at their N-terminus (Fig. 1B). A FLAG-tagged version of wild-type ZIKV NS5 (strain H/PF/2013), which localizes in the nucleus of infected Huh7 cells [12], was included in the analysis as a comparison. We also generated, based on previous data obtained with the ZIKV strain MR766 [12], a cytoplasmic mutant of ZIKV NS5 H/PF/2013 (referred to herein as NLSm), in which two positively charged residues (K390 and R393) of the NLS were mutated to alanine. 293T cells, which are easy to transfect, were used to assess the intracellular distribution of wild-type and mutant NS5 proteins of TBEV and ZIKV. Western blots analysis revealed that all NS5 TBEV mutants were expressed at their expected molecular weight but at a slightly lower levels than their wild-type counterparts (Fig. 1C). Similarly, reduced expression of the NS5 NLSm of ZIKV strain MR766, as compared to ZIKV NS5 wild-type, was previously observed in transfected 293T cells [11]. These data, together with ours (Fig. 1C), suggest that the importin-binding sites could be involved in NS5 stability. In line with previous microscopic analysis performed in 293T cells [12], cell fractionation experiments revealed that the majority of ZIKV NS5 wild-type was found in the nuclear fraction when expressed on its own. In contrast, the ZIKV NS5 NLSm was barely detected in the nuclear fraction (Fig. 1D). As in HMC3 cells infected with TBEV for 24 hours (Fig. 1A), similar amounts of exogenously expressed wild-type TBEV NS5 protein were found in the cytoplasmic and nuclear fractions of 293T cells (Fig. 1D). Deletion of β1NLS, α/βNLS, or both domains, show similar amount of NS5 into the cytoplasm, but reduced the amount of NS5 in the nucleus (Fig. 1D), suggesting that these 2 domains are involved in TBEV NS5 nuclear import. To identify the critical residues that are required for TBEV NS5 nuclear localization, all the basic residues within the two importin-binding domains were mutated to alanine. They were mutated either individually, in pairs, or in threes when near each other (Fig. S1D). Expression levels of the mutants were similar, suggesting that point mutations had little impact on the protein stability (Fig. S1E). All single and double mutants, like wild-type NS5, were equally distributed between the cytoplasmic and nuclear fractions (Fig. S1F). By contrast, the triple mutant K389A+K391A+R393A, which we named NLS mutant (NLSm), was predominantly in the cytoplasmic fraction (Fig. S1F and Fig. 1E). Quantification of epifluorescence microscopic images performed in HMC3 cells confirmed that the Δα/β deletion mutant and the NLSm exhibited a significantly reduced nuclear localization (Fig. 1F-G). Thus, the three positively charged residues of the α/βNLS, which are predicted to be surface-exposed (Fig. 1H), appeared to be key for NS5 nuclear localization.

To explore the importance of the NS5 NLS for TBEV replication, we took advantage of a replicon derived from the Neudoerfl strain [20], in which we mutated the 3 critical positively charged residues of the α/βNLS to alanine (replicon NLSm). We included in the analysis a wild-type replicon and a replication_defective replicon bearing double D_to_A mutations in the RdRp catalytic sequence GDD (replicon_GAA) [9], as positive and negative controls, respectively. *In vitro* transcripts synthesized from plasmids coding for these three replicons were electroporated into HMC3 cells and viral RNA levels were measured by RT-qPCR analysis at various time points post-electroporation. As expected, the wild-type replicon produced increasing amount of viral RNA over time, whereas the replicon_GAA did not (Fig. 1I). RNAs derived from the NSLm replicon were unable to replicate (Fig. 1I), indicating that the three NLS residues, and potentially nuclear localization of NS5, are critical for viral replication.

### The TBEV NS5 interactome is enriched in nuclear proteins

To identify cellular partners of TBEV NS5 during infection, we used IP-MS in 293T and HMC3 cells infected for 24 hours with the Hypr strain at an MOI of 1 (Fig. 2A). In these conditions, approximately 90% of cells were positive for the viral protein E at 24 hours post-infection (hpi) (Fig. 2B). TBEV NS5 was purified from infected cells using an in-house NS5 antibody. Non-infected cells were used as negative controls. The eluates were first analyzed by silver staining (Fig. S2A) and then by data-independent acquisition mass spectrometry (DIA-MS). Over 1,900 host proteins were detected in the purified samples from both cell lines (Fig. S2B). NS5 protein intensities in infected cells exceeded over 256-fold those of mock-infected controls (Fig. S2C). Pairwise correlation analysis confirmed the reproducibility of protein intensity profiles across biological replicates (Fig. S2D). We further analyzed the data with SAINTq, an interaction scoring algorithm optimized for DIA-MS data [21]. A total of 677 proteins, that had a Bayesian False Discovery Rate (BFDR) < 0.05, were considered potential partners of TBEV NS5 (Table S1).

**Figure 2.**
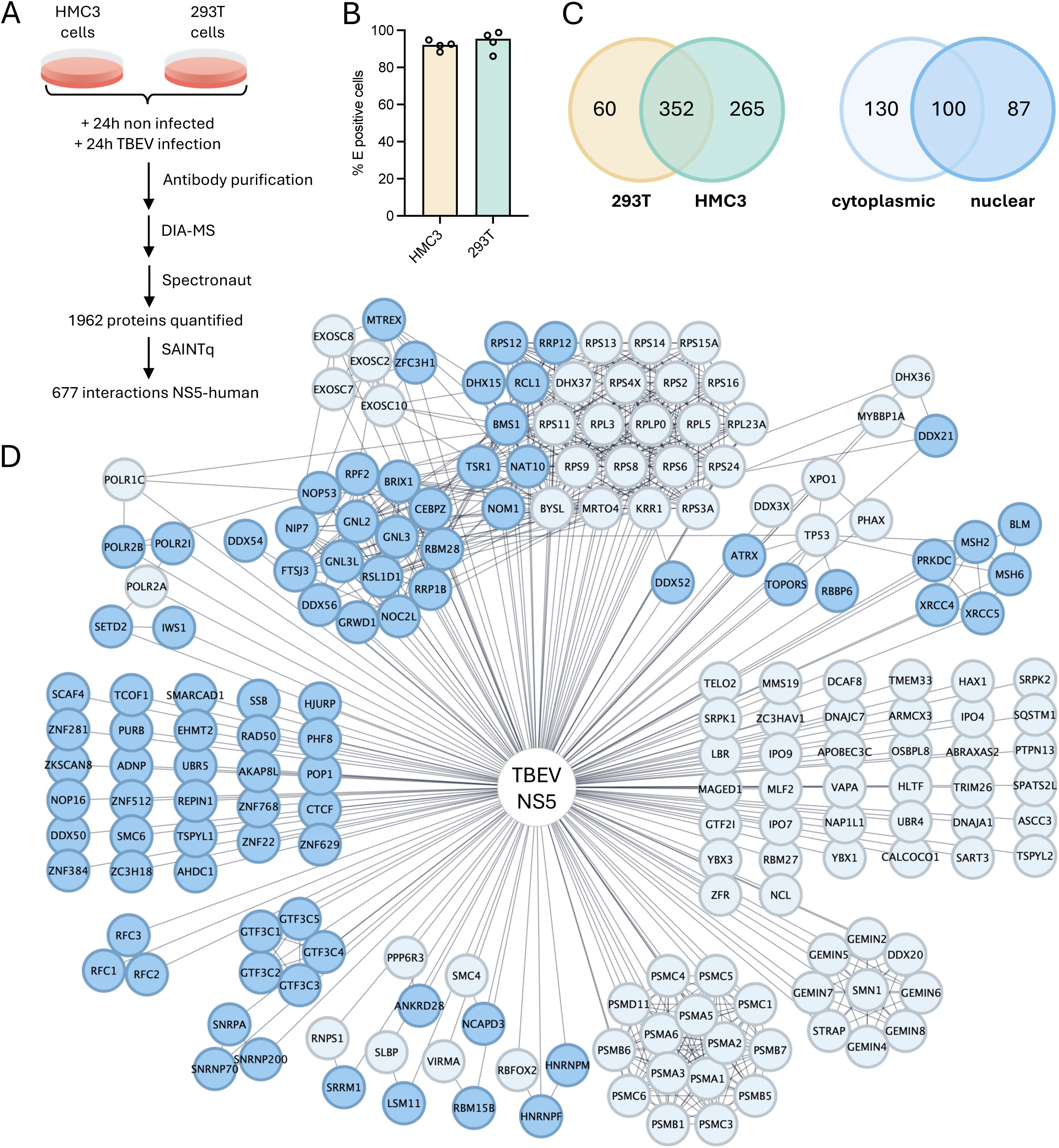
Identification of NS5 partners during TBEV infection. A. Experimental flow. HMC3 and 293T cells were mock-infected or infected with TBEV. After 24 hours, cell lysates were immunoprecipitated with anti-NS5 coupled to magnetic beads. Immunoprecipitated samples were analyzed by mass spectrometry (MS) using four independent biological replicates. Proteomic data was analyzed using SAINTq for interaction scoring. B. The percentage of infected cells at 24 hours post-infection was assessed by flow cytometry analysis using antibodies against the viral E protein. C. Venn diagrams of TBEV NS5-interacting partners identified by MS. The left panel shows the overlap of NS5 partners between the two cell lines. The right panel depicts the subcellular localization of 327 partners (out of the 352 identified in both cell lines), as determined by UniProt protein ID annotations. Of note, 25 proteins had unknown subcellular localization. D. Nuclear partners of TBEV NS5 identified in both HMC3 and 293T cells are represented. Partners with exclusive nuclear localization are depicted in dark blue, while those with partial nuclear localization are depicted in light blue. Protein-protein interaction networks were mapped using STRING analysis.

Among these partners, 352 proteins were common to both cell lines (Fig. 2C). Sixteen of these were recently identified as NS5 TBEV partners in an MS analysis performed in transfected 293T cells [22], such as SETD2 and NKTR. Protein ID inquiries using UniProt revealed that most of these 352 partners localizes exclusively or partially to the nucleus (Fig. 2C), consistent with the nuclear localization of NS5 (Fig. 1). Gene Ontology analysis of the TBEV NS5 nuclear partners showed that they participate in chromosome organization and remodeling, as well as RNA splicing, methylation, and metabolism, such as the members of the Gemin-SNM complex [23] (Fig. 2D and S3). This suggests that TBEV NS5 may modulate host epigenetic, transcriptomic, and epitranscriptomic programs to facilitate viral replication. Additionally, numerous ribosomal proteins, as well as about 15 proteasomal proteins, were identified, suggesting that NS5 may also associate with these complexes (Fig. 2D). Several NS5 interactors, such as ATRX and ZRANB2, were previously identified by MS analyses as partners of ZIKV, DENV and WNV NS5 [16,17], indicating that these proteins are conserved partners of orthoflavivirus NS5.

To assess the conservation of these interactions across tick-borne orthoflaviviruses, we compared our TBEV NS5 interactome with that of POWV NS5, which was identified through affinity purification (AP) MS in 293T and HMC3 cells expressing Strep-tagged NS5 proteins of 2 POWV strains (LB and SPO) (manuscript in preparation). Seventeen proteins were shared across at least 4 of the 6 datasets (Fig. 3A, top panel). Since these candidates were identified in two cell lines (293T cells and HMC3), either during infection for TBEV or in cells expressing POWV NS5, they were considered high confidence partners of tick-borne orthoflavivirus NS5. Among these, SCRIB and TYK2 were previously identified by us and others as direct partners of TBEV NS5 in a variety of cellular models, including yeasts and human cell lines [9,24,25], further validating our approach. We were also interested in potential TBEV NS5-specific partners. Three nuclear proteins (PHAX, HAX1 and SMRCD), which emerged as top candidates based on their BFDR scores in 293T and HMC3 cells infected with TBEV (Fig. 3A, top panel), were not identified as POWV NS5 partners. These 3 candidates, along with the seventeen shared proteins (excluding TYK2, whose role in viral replication is already well characterized [9]), were selected for further functional investigations.

**Figure 3.**
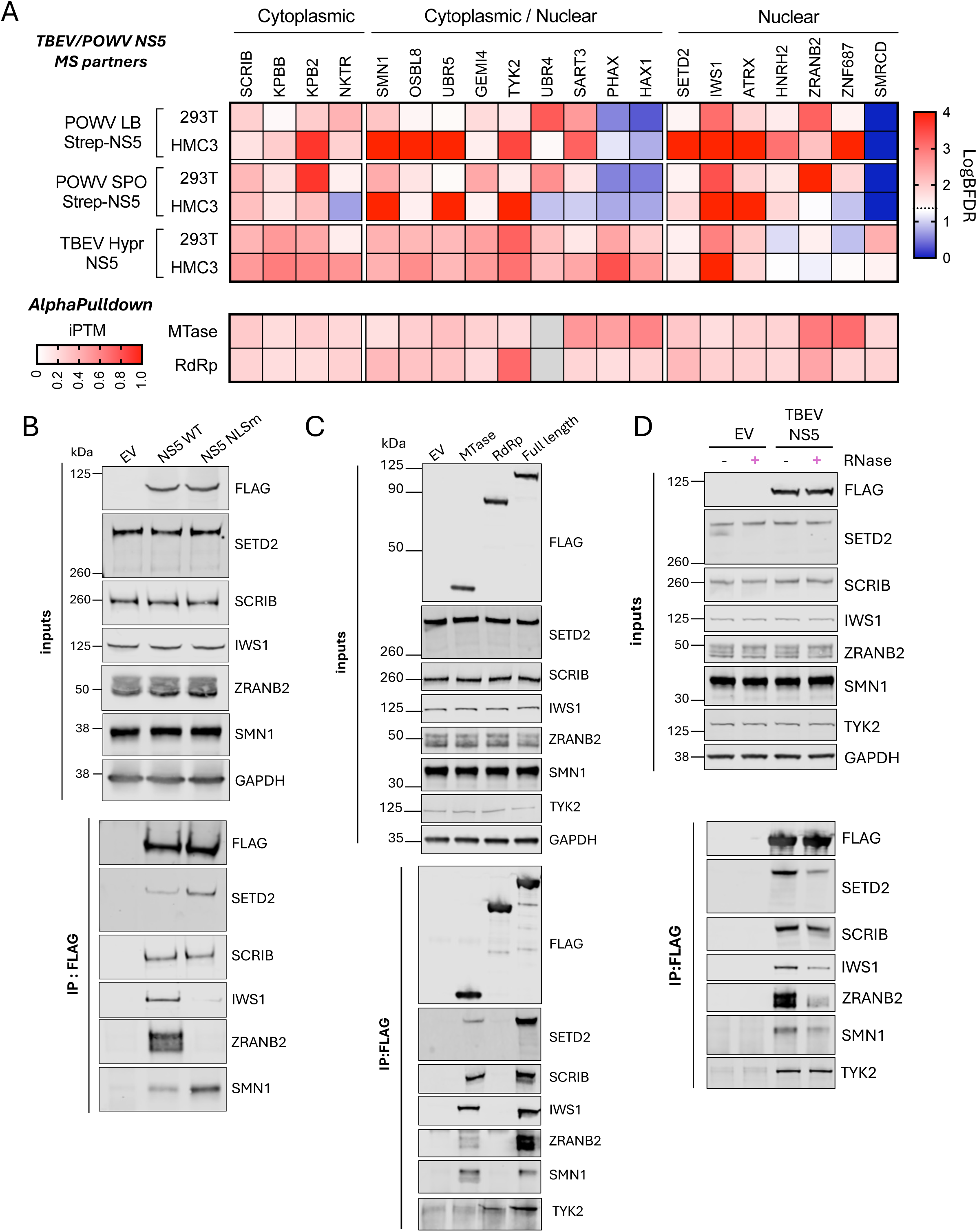
Validation of NS5 interaction with high-confidence nuclear and cytoplasmic partners. A. Upper panel: heat map representation of LogBFDR scores of NS5-interacting proteins identified by MS for TBEV and POWV NS5. Proteins with LogBFDR > 1.3 in at least 4 of the 6 datasets are displayed and considered as high-confidence NS5 partners. Lower panel: heat map of iPTM scores from an AlphaPulldown screen between TBEV-NS5 MTase or RdRp domains and the top 20 host protein partners. The iPTM value for UBR4 was not computed, as the protein siez exceeded computationally allowed limit. B. 293T cells were transfected for 24 h with plasmids encoding FLAG-tagged WT or NLSm NS5 TBEV proteins, or an empty vector as control (EV). Cell lysates were immunoprecipitated using FLAG antibodies. Western blot analysis was performed on whole cell lysates (inputs) and FLAG-immunoprecipitates (IP) using the indicated antibodies. C. 293T cells were transfected for 24 h with plasmids encoding FLAG-tagged MTase, RdRp or full length NS5 TBEV proteins, or an empty vector as control (EV). Cell lysates were immunoprecipitated using FLAG antibodies. Western blot analysis was performed on whole cell lysates (inputs) and FLAG-immunoprecipitates (IP) using the indicated antibodies. D. 293T cells were transfected for 24 h with plasmids encoding FLAG-tagged NS5 TBEV protein, or an empty vector as control (EV). Cell lysates were treated with RNase A or left untreated, and were immunoprecipitated using FLAG antibodies. Western blot analyses were performed on whole cell lysates (inputs) and FLAG-immunoprecipitates (IP) using the indicated antibodies.

To validate some of the interactions identified through MS, we performed co-immunoprecipitation assays in 293T cells expressing FLAG-tagged wild-type or NLSm TBEV NS5. To ensure comparable expression levels between the NS5 versions (Fig. 1), cells transfected with NS5 NLSm received four times the amount of plasmid used for wild-type NS5. We selected five high confident partners based on the availability of commercial antibodies: ZRANB2, IWS1, SETD2, SMN1 and SCRIB. ZRANB2 functions as a splicing factor [26], while IWS1 is a mRNA splicing and export factor that recruits SETD2 to chromatin to promote histone methylation [27]. SETD2 has also been reported to mediates STAT1 methylation in the cytoplasm [28]. SMN1 localizes to both the cytoplasm and the nucleus, where it is found in Gems and Cajal bodies [29], whereas SCRIB is primarily found at the cell surface [30]. Endogenous SETD2, SCRIB, IWS1, ZRANB2 and SMN1 co-precipitated with wild-type TBEV NS5 in transfected cells (Fig. 3B), confirming the interaction identified by IP-MS in infected cells (Fig. 2D). ZRANB2 and IWS1 showed no or minimal precipitation with the NLSm version of NS5 (Fig. 3B), indicating that these interactions are predominantly nuclear. This aligns with ZRANB2 and IWS1 localization, which is mainly nuclear in both non-infected and infected HMC3 cells (Fig. S4A), as previously described in other cellular models [25-26]. In contrast, SETD2, SMN1 and SCRIB co-precipitated with NS5 NLSm, suggesting these interactions occur, at least in part, in the cytoplasm (Fig. 3B). This is consistent with SMN1 localizing in the cytoplasm and nucleus of infected and non-infected HMC3 cells (Fig. S4A), as well as in human lung fibroblast MRC5 cells [29]. These findings also align with the detection of SCRIB at the plasma membrane of HMC3 cells (Fig. S4A), as well as with interactions between SCRIB and a V5-tagged TBEV NS5 at the surface of transfected canine kidney cells [24].

To predict whether the 20 selected putative NS5 TBEV partners preferentially bind the RdRp or MTase domain, we used Alphapulldown [31,32]. The interactions between the MTase domain and HAX1, ZRANB2, and ZNF687 yielded interface predicted template modelling (iPTM) scores above 0.6, suggesting potential direct interactions (Fig. 3A, lower panel, Table S2). The iPTM score for the interaction between the RdRp domain and TYK2 was the highest among all partners (0,74) (Table S2), consistent with yeast experiments demonstrating a direct interaction [9]. Co-immunoprecipitation assays in 293T cells expressing FLAG-tagged versions of the full-length NS5, the RdRp, or the MTase domains were performed. TYK2 served as a control [9]. SETD2, SCRIB, IWS1, ZRANB2 and SMN1 co-precipitated with the MTase domain (Fig. 3C). These results align with some of the Alphapulldown predictions (Fig. 3A, lower panel), supporting the domain-specific nature of these interactions.

To determine whether the interactions between NS5 and these 5 partners are mediated by RNA, cell extracts were treated with RNAse A prior to immunoprecipitation. As positive controls, we used cells expressing tagged versions of IGF2BP2 and IGF2BP3, two RNA-binding proteins known to form RNA-dependent heterodimers [33,34]. As expected [33,34], RNase A treatment disrupted their interactions (Fig. S4B). Co-immunoprecipitation of SETD2, IWS1, ZRANB2, and SMN1 with NS5 was reduced in RNase A treated extracts, compared to untreated controls, while interactions with SCRIB and TYK2 remained unaffected (Fig. 3D). The resistance of SCRIB and TYK2 interactions to RNase A treatment is consistent with previous reports of direct protein-protein interactions between NS5 and TYK2 in yeast assays [9,25] and NS5 and SCRIB in GST pull-down experiments [24]. The interaction between NS5 and ZRANB2 was particularly sensitive to RNAse A treatment, suggesting that these 2 proteins interact in an RNA-dependent manner. Since ZRANB2 remained localized in the nucleus during viral infection (Fig. S4A) and interact with NS5 in the nucleus (Fig. 3B), its binding to NS5 is likely mediated by cellular RNAs.

Together, these results showed that NS5 engages in both RNA-dependent and direct protein-protein interactions with host factors, some occurring in the nucleus and others in the cytoplasm.

### Reduced expression of some high-confidence TBEV NS5 partners modulates viral replication

To assess whether the nineteen high-confidence TBEV NS5 partners that we selected (Fig. 3A) impact viral replication, a loss-of-function screen was performed (Fig. 4A). HMC3 cells were transfected with pools of four siRNAs targeting the selected NS5 interactants and infected 48_h later with TBEV at an MOI of 0.001. siRNAs targeting ATP6V0C, a protein promoting fusion between the endosomal membrane and the envelope of orthoflaviviruses [35] were used as positive controls. Three sets of non-targeting siRNA pools were used as negative controls. At 48_h post-infection, intracellular viral RNA levels were evaluated by RT-qPCR analysis (Fig. 4B) and the number of cells positive for the viral protein E were quantified by flow cytometry analysis (Fig. 4C). Supernatants were also harvested and incubated on Vero cells to evaluate infectious viral particle production (Fig. 4D). cDNA samples were used to analyze mRNA levels of the selected genes and assess knock-down efficiency (Fig. 4E). Finally, cell viability was measured at 72_h post-transfection in mock-infected samples (Fig. 4F). Viral replication measurements were compared to those obtained from cells transfected with the non-targeting siRNA pool. As expected [35], reducing ATP6V0C significantly decreased viral RNA yield, viral proteins expression and viral particles production without affecting cell viability (Fig. 4B-F). Reduced expression of eight genes (UBR5, SCRIB, HNRNPH2, ZNF687, GEMIN4, SMN1, SETD2 and SMARCAD1) decreased viral RNA yield, the number of infected cells and viral titers by over 50% compared with cells transfected with control siRNAs (Fig. 4B-D). Reduced expression of nine NS5 partners had little to no effect on TBEV replication (Fig. 4B-D). ZRANB2 and SART3 behaved like antiviral gene since knocking down their expression significantly increased viral RNA yield and the percentage of infected cells (Fig. 4B-D). RT-qPCR analyses revealed that all the siRNA pools were reducing the expression of their respective targets by at least 50% (Fig. 4E). The viability of cells treated with siRNAs against UBR5 and SART3 decreased by over 20% compared to control cells (Fig. 4F). Therefore, data related to UBR5 and SART3 silencing should be interpreted with caution. Seven of the eight NS5 partners that significantly modulated viral replication have nuclear activities (HNRNPH2, ZNF687, GEMIN4, SMN1, SETD2, SMARCAD1 and ZRANB2), further suggesting that nuclear NS5 plays a role in the viral life cycle.

**Figure 4.**
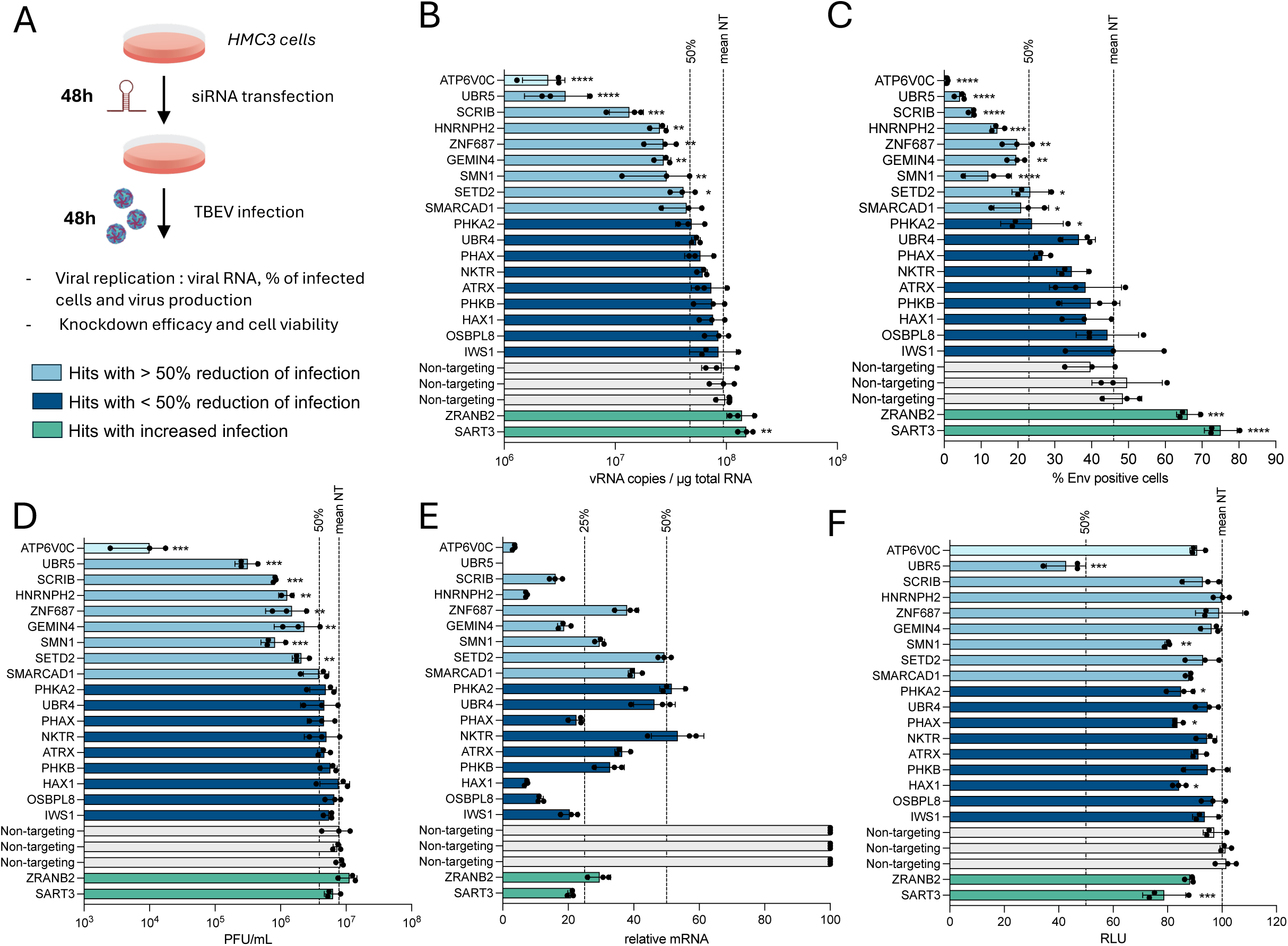
Functional screening of NS5 high-confidence interactants during TBEV infection. A. Scheme summarizing the screen conditions. HMC3 cells were transfected with either a pool of siRNAs against the indicated candidate genes or non-targeting (NT) siRNAs and infected with TBEV (at an MOI of 0.001 PFU/cell) for 48 hours. B. Cell-associated viral RNA was quantified by RT-qPCR and expressed as vRNA copies / µg of total RNA. C. The percentage of infected cells was assessed by flow cytometry analysis using antibodies against the viral E protein. D. The amount of released infectious virus particles in the supernatants of infected cells was quantified by plaque assays. E. Efficacy of the specific siRNAs. The relative abundances of the mRNAs of the candidate genes were determined by RT-qPCR analysis and were normalized to GAPDH mRNA levels. Values are expressed relatively to abundance in cells transfected with NT siRNAs in each experiment, set as 100. F. Cell viability of siRNAs transfected cells was assessed using CellTiter-Glo Luminescent Cell Viability Assays. Data are means ± SD of three independent experiments, *p < 0.05, **p < 0.01,***p < 0.001, ****p < 0.001, one-way anova test.

## Discussion

Since NS5 is too large to passively diffuse between the nucleus and the cytoplasm, it relies on active transport through the nuclear pore for nuclear import. We identified three residues, K389, K391 and R393, within the α/βNLS, that are essential for the nuclear import of TBEV NS5. Similar findings have been reported for mosquito-borne orthoflaviviruses, where K390 and R393 in ZIKV, the REE motif (residues 396–398) in DENV2, and the REK motif (residues 389–393) in WNV play pivotal roles in NS5 nuclear localization [10,12,36]. Thus, the α/βNLS appear to be a conserved primary determinant of NS5 nuclear addressing across orthoflaviviruses. Notably, DENV2 also possesses a C-terminal NLS, indicating virus-specific features within the *orthoflavivirus* genus [37]. While deletion of the β1NLS domain impaired NS5 nuclear localization, mutation of its positively charged residues did not, suggesting that the deletion may induce conformational changes that indirectly disrupt nuclear import. Although we did not identify β-importin1 or α-importin proteins as cellular partners of TBEV NS5, we found IPO4, IPO7 and IPO9, which are members of the importin family, as specific interactors. Future studies could involve reducing the expression of these proteins to assess their role in NS5 nuclear import. We also identified Exportin-1 (XPO1) as a cellular partner of TBEV NS5, as previously shown for DENV NS5 [38], suggesting that NS5 actively shuttles between the cytoplasm and the nucleus. However, the export mechanisms of NS5 remain poorly understood and warrant further investigation.

Previous studies have suggested that shuttling between the cytoplasm and the nucleus protect ZIKV NS5 from degradation [11]. A similar mechanism may apply to TBEV, as NS5 NLSm was less expressed than wild-type NS5, though to a lesser extent than for ZIKV NS5. Mutation of key residues for NS5 nuclear import abolished viral RNA replication of a TBEV replicon. While these mutations do not lie within the catalytic pocket of the RdRp domain, they could induce conformational changes that indirectly impair replication of the NLSm. In the case of DENV, ZIKV and WNV, RdRp activity of the NS5 NLSm showed only marginally, if any, impact on the polymerase function [10,11,39], suggesting that defective replication is not due to a loss of enzymatic activity. However, the absence of RNA replication, even at very early stages, suggests that additional factors might be at play, like reduced interactions between NS5 and viral or host proteins required for replication. In DENV, viral replication was unaffected by NS5 mutants with reduced nuclear localization, suggesting that the extent of nuclear import is not directly correlated with replication efficiency [10]. Importin inhibitors used to block NS5 nuclear import in DENV- and ZIKV-infected cells significantly reduced infection [40–42]. However, their broad effect on the cells prevents definitive conclusions about the specific role of nuclear NS5 in viral replication. Assessing the significance of NS5 nuclear localization in an infectious context remains challenging, as both mutational and pharmacological inhibition approaches may introduce indirect effects.

Previous interatomic studies of orthoflavivirus proteins have relied on tagged versions of proteins expressed individually [16,17] or from modified infectious clones [14]. Here, we used IP-MS to identify NS5 partners in infected human cells. Comparing the NS5 partners we identified previously by immunoprecipitation of FLAG-tagged NS5 expressed in 293T cells [9] with those recovers here (by immunoprecipitating NS5 from infected 293T or HMC3 cells using NS5 antibodies) revealed limited overlap, with only 35 common partners. This divergence may be attributed to the use of different antibodies. Immunoprecipitation via the FLAG tag offers the advantage of better-controlled background noise, as associated contaminants are well-characterized. In contrast, the use of anti-NS5 antibodies, while enabling analysis in the context of infection, may introduce higher background noise. However, the inclusion of non-infected cells as controls helps mitigate this effect. Additionally, differences in the lysis buffers used in the two protocols may influence the profile of detected interactions, particularly impacting indirect interactions and multiprotein complexes. Among the 35 partners identified by both approaches are TYK2, ATRX, SMRCD and members of the GEMIN complex. These proteins were also identified as partners of POWV NS5 partners through AP-MS in 293T and HMC3 cells expressing Strep-tagged NS5 proteins (manuscript in preparation).

We validated the interactions between SETD2, SCRIB, IWS1, ZRANB2 and SMN1, with NS5 by co-immunoprecipitation. The binding of SETD2, IWS1, and SMN1 to NS5 was reduced in transfected cells extracts treated with RNase A, suggesting that cellular RNA may stabilize these interactions. The interaction between NS5 and ZRANB2 was highly sensitive to RNase A treatment, indicating that these two proteins require RNA for their association. Indeed, the potential binding of NS5 to cellular RNAs has not been explored. To identify potential cellular RNAs bound to NS5, one approach could involve immunoprecipitation of NS5, either from transfected or infected cells, followed by RNA extraction and RNA-seq analysis.

The strong enrichment of nuclear proteins among NS5-associated cellular factors suggests that NS5 may have multiple functions in the nucleus. We identified six NS5 partners with described nuclear activities (HNRNPH2, ZNF687, GEMIN4, SMN1, SETD2 and SMARCAD1) that act as proviral factors for TBEV replication. GEMIN4, SMN1, HNRNPH2 and ZNF687 are all involved in splicing processes, while SETD2 and SMARCAD1 participate in epigenetic regulation. Similar to DENV NS5, TBEV NS5 may co-opt these proteins to modulate host splicing and gene expression, thereby suppressing antiviral responses and promoting replication [14,17,43]. Interestingly, the interaction of the cytoplasmic NS5 mutant with SETD2 and SMN1 suggests that these interactions occur, at least partly, in the cytoplasm. This raises the possibility that NS5 alters the subcellular distribution of these proteins, preventing them from performing their nuclear function. However, we did not observe a relocalization of these proteins during infection.

Finally, we identified ZRANB2 as an antiviral nuclear partner of TBEV NS5. ZRANB2 is an RNA-binding protein involved in pre-mRNA splicing [26,44,45]. It also interacts with the NS5 protein of WNV [16], DENV, ZIKV [17] and JEV [15] and inhibits the replication of WNV, DENV, and ZIKV [16]. Interestingly, its phosphorylation level increases during DENV and WNV infections [46,47]. These findings suggest that ZRANB2 is a conserved nuclear partner of orthoflaviviruses NS5 and may play an antiviral role in their replication. Our data showed that it interacts with the MTAse domain of TBEV NS5, predominantly in the nucleus of infected cells, and in an RNA-dependent manner. Further characterization of the interaction between NS5 and ZRANB2 could help determine its function during viral replication. Additionally, analyzing the spliceosome composition in infected cells, with or without ZRANB2, may reveal changes in the expression of host factors critical for viral replication.

Importantly, some cellular partners that did not appear to modulate viral infection in our functional assay may still play critical roles in the viral life cycle. This limitation arises because our assays were constrained by several factors, including the use of a single cell type, a unique time point, and an *in vitro* environment that does not fully recapitulate the complexity of viral infection *in vivo*. Further investigations into the nuclear activities of TBEV NS5 are therefore essential.

## Methods

### Cells and viruses

Human embryonic kidney (HEK) 293T cells and Human Microglial Clone 3 (HMC3) cells were obtained from the American Type Culture Collections (ATCC, CRL-3126 and CRL-3304). Huh7 human hepatocellular carcinoma cells were kindly provided by A. Martin, Institut Pasteur Paris. African green monkey kidney epithelial Vero cells were kindly provided by F. Tangy, Institut Pasteur Paris. All cultures were grown under standard conditions in Dulbecco’s Modified Eagle Medium containing GlutaMAX (Gibco) and supplemented with 10% heat-inactivated fetal bovine serum (FBS), and 1% penicillin and streptomycin (10,000 IU/mL; Thermo Fisher Scientific).

Experiments with TBEV were performed in a BSL-3 laboratory, following safety and security protocols approved by the risk prevention service of Institut Pasteur. The TBEV strain (Hypr strain, isolated in Czech Republic in 1953) was obtained from the European Virus Archive (EVAg, 008v-EVA285). Virus stocks were produced on Vero cells. Titration of infectious virions was performed by plaque assay on Vero cells, as previously described for other orthoflaviviruses [48]. HMC3 and 293T cells were infected at the MOI indicated in the figure legends, for 2 hours in DMEM medium containing 2% FBS, followed by replacement with 10% FBS media.

### Antibodies

Anti-TBEV NS5 antibodies were obtained by immunization of a rabbit using full-length recombinant TBEV NS5, as previously described [9]. They were used at a dilution of 1:1,000 for western blot and immunofluorescence assays. Anti-Envelope Mab 4G2 pan-orthoflavivirus antibody (kind gift from Philippe Desprès) was used at a dilution of 1:1,000 for flow cytometry and immunofluorescence assays. Anti-FLAG M2 (F3165, Sigma Aldrich) was used at a dilution of 1:5,000 for western blot. Anti-lamin B1 (12987-1-AP, Proteintech) and anti-GAPDH (10494-1-AP, Proteintech) were used at a dilution of 1:5,000 and 1:10,000 for western blot, respectively. Anti-IWS1 (#5681, Cell Signaling Technology), anti-SETD2 (#80290, Cell Signaling Technology), anti-Scribble (#4475, Cell Signaling Technology), anti-SMN1 (#12976, Cell Signaling Technology) and anti-ZRANB2 (24816-1-AP, Proteintech) and anti-TYK2 antibodies [9] were used at a dilution of 1:1,000 for western blot and 1:200 for immunofluorescence. The following secondary antibodies were used: Alexa Fluor 488 goat anti-mouse IgG (H+L) (A11001, Invitrogen); Alexa Fluor 647 donkey anti-mouse IgG (H+L) (A31571, Invitrogen) Alexa Fluor 647 goat anti-rabbit IgG (H+L) (A21244, Life Technologies); Alexa Fluor 680 goat anti-mouse IgG (H+L) (A21058, Invitrogen) and goat anti-rabbit IgG (H+L) Dylight 800 (SA5-35571, Invitrogen). All secondary antibodies were used at a dilution of 1:1,000 for flow cytometry and immunofluorescence assays and 1:10,000 for western blot analysis.

### Plasmids and cloning

Plasmids coding for full-length, MTase and RdRp domains of TBEV NS5, as well as ZIKV NS5, in p3xFLAG-CMV10 (Sigma) were previously described [9]. Deletion mutants of TBEV NS5 were generated by PCR using p3xFLAG-WT-TBEV-NS5 as a template and the primers listed in Table S3. Point mutants of TBEV and ZIKV NS5 were generated by site-directed mutagenesis of p3xFLAG-WT-NS5 or TBEV replicon using the primers listed in Table S3 and the Phusion Site-Directed Mutagenesis Kit (10493438, Fisher Scientific), following manufacturer’s instructions. Plasmids encoding IGF2BP1 and IGF2BP3 were from Eugene Yeo’s and Thomas Tuschl’ laboratories, respectively, and were acquired through Addgene (#155676 and #19879). All plasmids were amplified in TOP10 cells (Thermo Fisher Scientific) and sequencing was outsourced to Eurofins genomics.

### TBEV replicons

WT and GAA replication defective TBEV replicons were previously described [9]. TBEV pTND/ΔME replicon plasmids [20] were linearized by NheI digestion and blunt ended using Quick Blunting Kit (New England Biolab). Five µg of purified DNA template were used for T7 *in vitro* transcription using RiboMAX large-scale RNA production system T7 (Promega) in presence of 40 mM cap analog (Ribo m7G Cap, Promega) following the manufacturer’s instructions. After RQ1 DNase treatment (Promega), RNA was purified with RNA clean-up kit (Macherey-Nagel). *In vitro* synthetized TBEV replicon RNA was introduced into HMC3 cells by electroporation. Briefly, 2.5 × 10^6^ trypsinized HMC3 cells were washed tree time in cold PBS, resuspended in 400 µl cold PBS and electroporated with 2.5 µg of RNA in 0.4-cm electroporation cuvettes (Biorad) with a 25 µF and 450 V pulse using Genepulser system (Biorad). After electroporation, cells were collected in 12 ml of warm medium, cell suspension was transferred to 12 well plates (1 mL per well) and incubated at 37°C under standard conditions.

### Plasmid transfections

293T and HMC3 cells were transfected using TransIT-293 (Mirus) and FuGENE HD (Promega) transfection reagents, respectively, following the manufacturer’s protocol. One day later, samples were processed for western-blot and co-immunoprecipitation.

### Western blot

Cells were lysed in Radio-Immunoprecipitation Assay RIPA buffer (Sigma-Aldrich) supplemented with a protease inhibitor cocktail (Roche) for 15 min at 4°C and lysates were clarified by centrifugation at 10,000 x g at 4°C for 15 min. Samples were denatured at 95°C in 1x Protein Sample Loading Buffer (Li-Cor Bioscience) under reducing conditions (NuPAGE reducing agent, Thermo Fisher Scientific). Proteins were separated by SDS-PAGE using NuPage 4-12% Bis-Tris gels (Invitrogen) and transferred to nitrocellulose membranes (Bio-Rad) using a Trans-Blot Turbo Transfer system (Bio-Rad). Alternatively, high molecular weight proteins were separated by SDS-PAGE using NuPAGE 3-8% Tris-Acetate gels (Invitrogen) and were transferred to nitrocellulose membranes using liquid transfer buffer (50 mM Tris, 25 mM glycine, 0.4% SDS, 20% ethanol) for 2 hours at 25 V. Membranes were blocked with PBS-0.1% Tween 20 (PBS-T) containing 5% milk and incubated overnight at 4°C with primary antibodies diluted in blocking buffer. Finally, membranes were washed and incubated for 45 min at room temperature with secondary antibodies diluted in blocking buffer and washed again. Images were acquired using an Odyssey CLx infrared imaging system (Li-Cor Bioscience).

### Co-immunoprecipitation

293T cells were transfected with FLAG-tagged NS5 plasmids for 24 hours and were lysed in IP lysis buffer (50 mM Tris-HCl pH 7.5, 150 mM NaCl, 1 mM EDTA, Complete protease inhibitor tablet (EDTA-free) and 0.5% IGEPAL CA-630) for 30 min at 4°C. Lysates were clarified by centrifugation at 5,000g for 20 min at 4°C. Clarified lysates were incubated with magnetic beads coupled with anti-FLAG M2 (M8823; Sigma-Aldrich) or anti-V5 (SAE0203; Sigma-Aldrich) overnight at 4°C. Following incubation, beads were washed four times with IP washing buffer (50 mM Tris-HCl pH 7.5, 150 mM NaCl, 1 mM EDTA, and 0.05% IGEPAL CA-630), then once with detergent-free IP buffer (50 mM Tris-HCl pH 7.5, 150 mM NaCl and1 mM EDTA), and proteins were eluted with 1X Protein Sample Loading Buffer (Li-Cor Bioscience) and 1X NuPAGE reducing agent (Thermo Fisher Scientific). For nuclease experiments, clarified lysates were treated with 100 µg/mL of RNAse A (EN0531; Thermo Fisher Scientific) for 20 min at room temperature prior to immunoprecipitation. Proteins of interest were detected by western blot, as described above.

### Cellular fractionation

The protocol was adapted from [49]. 293T cells were harvested and washed once with ice-cold PBS. After 5 min of centrifugation at 300 x g, cells were lysed with low salt lysis buffer (10 mM Tris pH 7.5, 10 mM NaCl, 3 mM MgCl2, 0.3% IGEPAL CA-630 and 10% glycerol) for 10 min on ice to preserve nucleus integrity. After centrifugation at 800 x g for 8 min at 4°C, the supernatant was saved for “cytoplasmic” fraction. The pellet containing the nuclei was washed 4 times with low salt lysis buffer by centrifugation at 300 x g for 4 min at 4°C. Finally, the pellets containing washed nuclei were lysed with a high salt denaturing buffer (10 mM Tris-HCl pH 7.0, 4 mM EDTA, 0.3 M NaCl, 1 M urea, and 1% IGEPAL CA-630) to generate nuclear extracts containing solubilized nuclear proteins and protein stably bound to the chromatin.

### Immunofluorescence assays

HMC3 and Huh7 cells were plated into 24-well plates on glass coverslips. After transfection or infection, wells were fixed with 4% paraformaldehyde (PFA) (Sigma-Aldrich) for 30 min at room temperature, permeabilized 50% v/v methanol/ethanol (Sigma-Aldrich) V/V for 15 min and then blocked for 30 min with PBS containing 0.05% Tween and 5% BSA before incubation with the indicated primary antibodies for 1 hour. After incubation, cells were washed three times with PBS containing 0.05% Tween. Secondary Alexa Fluor 488 or 647-conjugated antibodies and DAPI were added for 1hour, then cells were washed 3 times with PBS containing 0.05% Tween. Slides were mounted using Prolong gold (Life Technologies, P36930) imaging medium. Images were acquired using a Zeiss LSM700 confocal microscope, or a Zeiss Imager Z1 epifluorescence microscope. Quantification of the nuclear localization of NS5 WT or mutants was determined by quantifying intensity of cytoplasmic (C) and nuclear (N) FLAG signal using Fiji, followed by calculation of the ratio of mean fluorescence N/C.

### Flow cytometry

Infected cells were harvested using trypsin and fixed with cytofix/cytoperm kit (BD, Pharmingen). Cells were washed twice with Perm/Wash buffer (BD, Pharmingen) and stained using the anti-E Mab 4G2 primary antibody diluted in Perm/Wash buffer for 1 hour at 4°C. Cells were washed twice and stained with secondary anti-mouse Alexa 647 in the dark for 45 min at 4°C. After washing the cells twice, data were acquired using Attune NxT Acoustic Focusing Cytometer (Life Technologies) and analyzed using FlowJo software.

### RNA extraction and RT-qPCR analyses

Total RNAs from cells were extracted using the NucleoMag RNA kit (Macherey-Nagel) following the manufacturer’s protocol. First-strand cDNA synthesis was performed on 1 µg of total RNA with the RevertAid H Minus Moloney murine Leukemia Virus (M-MuLV) reverse transcriptase (Thermo Fisher Scientific) using random primers p(dN)_6_ (Roche). Quantitative real-time PCR was performed on a real-time PCR system (Quant Studio 6 Flex; Applied Biosystems) with SYBR green PCR master mix (Life Technologies). Data were analyzed by the ΔΔCt method, with all samples normalized to GAPDH. All experiments were performed in technical triplicate. The primers used for RT-qPCR are listed in Table S4. Quantification of TBEV genomes were determined by extrapolation from a standard curve generated from serial dilutions of the plasmid encoding TBEV NS5 (p3xFLAG-NS5-TBEV).

### Sample preparation for mass spectrometry (MS)

HMC3 and 293T cells were infected with TBEV for 24 hours, then harvested by trypsinization and counted. 500,000 cells were used for flow cytometry staining of viral envelope to check efficacy of infection. Non-infected cells were used as controls. For affinity purification of NS5, 10 million cells were lysed in 1 ml of IP lysis buffer (50 mM Tris-HCl pH 7.5, 150 mM NaCl, 1 mM EDTA, Complete protease inhibitor tablet (EDTA-free) and 0.5% IGEPAL CA-630) for 30 min at 4°C. Lysates were clarified by centrifugation at 5,000g for 20 min at 4°C. Clarified lysates were incubated with 10 µg of anti-TBEV NS5 antibody overnight at 4°C. Following incubation, 40 µl of Pierce Protein A/G magnetic beads (#88802, Thermo Fisher Scientific) were added to lysates – antibody mixture, and incubated for 2 hours at 4°C. Beads were washed four times with IP washing buffer (50 mM Tris-HCl pH 7.5, 150 mM NaCl, 1 mM EDTA, and 0.05% IGEPAL CA-630). 3 µL of beads were eluted with 1X Protein Sample Loading Buffer (Li-Cor Bioscience) and 1X NuPAGE reducing agent (Thermo Fisher Scientific). The eluate was analyzed by SDS PAGE and silver staining (Pierce). For MS, immunoprecipitates were directly digested on beads. Beads were washed once and resuspended with 40 µl of 2M urea, 50 mM Tris pH 8.0 and 1 mM DTT, and incubated for 30 min at 37°C with shaking at 700 rpm. Iodoacetamide (Sigma-Aldrich) was added to 3 mM final concentration and incubated at room temperature for 45 min in the dark with shaking. Digestion was performed by adding 3 mM DTT and 750 ng of sequencing grade modified trypsin (Promega) and incubating at 37°C with shaking overnight. Finally, supernatants were collected to fresh tubes and an additional 500 ng of sequencing grade modified trypsin was added and incubated for 2 hours at 37°C with shaking. Digested samples were desalted using C18 solid phase extraction columns (C18 BioPurSPN, Nest Group) according to the manufacturer’s instructions. Desalted samples were vacuum centrifuged to dryness and resuspended in 0.1% formic acid for mass spectrometry analysis.

### Mass spectrometry (MS) and data analyses

All samples were analyzed on an Orbitrap Eclipse mass spectrometry system equipped with an Easy nLC 1200 ultra-high pressure liquid chromatography system interfaced via a Nanospray Flex nanoelectrospray source (Thermo Fisher Scientific). Samples were injected onto a fritted fused silica capillary (30 cm × 75 μm inner diameter with a 15 μm tip, CoAnn Technologies) packed with ReprosilPur C18-AQ 1.9 μm particles (Dr. Maisch GmbH). Buffer A consisted of 0.1% formic acid in water, and buffer B consisted of 0.1% formic acid in 80% acetonitrile. Peptides were separated by an organic gradient from 5% to 35% mobile buffer B over 60 min, followed by an increase to 100% B over 10 min at a flow rate of 300 nL/min. Analytical columns were equilibrated with 3 μL of buffer A.

To build a spectral library, samples from each set of biological replicates were pooled and acquired in data-dependent manner. Data-dependent acquisition (DDA) was performed by acquiring a full scan over a m/z range of 375-1025 in the Orbitrap at 120,000 resolving power (@ 200 m/z) with a normalized AGC target of 100%, an RF lens setting of 30%, and an instrument-controlled ion injection time. Dynamic exclusion was set to 30 seconds, with a 10 p.p.m. exclusion width setting. Peptides with charge states 2-6 were selected for MS/MS interrogation using higher energy collisional dissociation (HCD) with a normalized HCD collision energy of 28%, with 3 seconds of MS/MS scans per cycle. Data-independent analysis (DIA) was performed on all individual samples. A full scan was collected at 60,000 resolving power over a scan range of 390-1010 m/z, an instrument controlled AGC target, an RF lens setting of 30%, and an instrument controlled maximum injection time, followed by DIA scans using 8 m/z isolation windows over 400-1000 m/z at a normalized HCD collision energy of 28%.

The Spectronaut algorithm was used to build spectral libraries from DDA data, identify peptides/proteins, and extract intensity information from DIA data [50]. DDA data were searched against the *Homo sapiens* UniProt reference proteome (downloaded on August 23, 2023) and TBEV protein sequences. False discovery rates were estimated using a decoy database strategy. All data were filtered to achieve a false discovery rate of 0.01 for peptide-spectrum matches, peptide identifications, and protein identifications. Search parameters included a fixed modification for carbamidomethyl cysteine and variable modifications for N-terminal protein acetylation, and methionine oxidation. All other search parameters were Biognosys factory defaults.

Statistical analysis of proteomics data was conducted utilizing the MSstats package in R [51] and SAINTq for interaction scoring [52]. For MSstats analysis, all data were normalized by equalizing median intensities, the summary method was Tukey’s median polish, and the maximum quantile for deciding censored missing values was 0.999. Interaction network of TBEV NS5 protein was represented using Cytoscape software (v3.9.1). Host protein-protein interactions and Gene Ontology analysis were performed using stingApp on cytoscape (https://apps.cytoscape.org/apps/stringapp).

### siRNA screening

Pools of 4 siRNAs for each selected host target (Horizon Discovery) were transfected in HMC3 cells using HiPerFect (Qiagen) transfection reagent according to manufacturer’s recommendations at a final siRNA concentration of 10 nM. 48 hours after transfection, cells were infected with TBEV at an MOI of 0. 001. At 48 hours post-infection, viral replication was assessed by measuring percentage of infected cells by flow cytometry, cell-associated viral RNA by RT-qPCR and amount of released infectious virus particles in the supernatants by plaque assays. Cytotoxicity of siRNAs was assessed at 72 hours post transfection using CellTiter-Glo Luminescent Cell Viability Assay (Promega) according to manufacturer’s instructions.

## Computational interaction analysis using AlphaFold 3

Computational predictions of protein–protein interactions were generated using AlphaFold 3 (AF3) [32]. All candidate sequences were retrieved from UniProt, and for TBEV-NS5, we used residues 1–255 for the MTase domain and 276-875 for the RdRp domain. AF3 was run with default parameters for multimolecular complex prediction on the AlphaFold Server (https://alphafoldserver.com), modelling each complex as a single 1:1 assembly. For each interaction, the average of five models was used, selected according to AF3 confidence metrics, including per-residue confidence and interface confidence (iPTM).

## Data representation and statistical analyses

Data are presented and analyzed using GraphPad Prism 10.

## Acknowledgements

This work was supported by the Institut Pasteur, Centre National de la Recherche Scientifique and Agence Nationale de la recherche (ANR-21-CE15-0031) awarded to NJ, as well as by NIH/NIAID grant numbers R01AI124690 and R01AI143850, awarded to J.R.J and H.R, respectively.

## Supplementary figure legends

**Figure S1. Nuclear localization of TBEV NS5.** A. Staining of NS5 (green) and E (red) of TBEV in mock-infected HMC3 cells. Nuclei were stained using DAPI (blue) and images were acquired using a confocal microscope. Scale bars 20 µm. B. Huh7 cells were infected with TBEV at an MOI of 1. After 24 hours, subcellular localization of NS5 was determined by immunofluorescence microscopy using antibodies against TBEV NS5 (green). Nuclei were stained using DAPI (blue) and images were acquired using a confocal microscope. Scale bars 20 µm. C. Sequence alignment of NLS importin β1- and α/β-interacting domains of several orthoflavivirus NS5 proteins. Positively charged residues are highlighted in red and arrows indicate conserved ones. D. Representation of TBEV NS5 point mutants generated by mutagenesis. Positively charged residues (in red) were mutated to alanine. E. 293T cells were transfected with plasmids encoding TBEV FLAG-NS5 WT or mutants. After 24 hours, whole-cell lysates were analyzed by western blot using antibodies against FLAG and GAPDH. F. 293T cells were transfected with plasmids encoding TBEV FLAG-NS5 WT or mutants. After 24 hours, NS5 subcellular localization was determined by analyzing cytoplasmic (C) and nuclear (N) fractions by western blot using antibodies against FLAG, GAPDH and Lamin B1.

**Figure S2. Mass spectrometry complementary data.** A. Representative silver staining of immunoprecipitated samples from HMC3 and 293T infected cells. Silver-stained bands corresponding to NS5 are indicated by arrows. B. Number of unique proteins quantified for each sample. C. Box plot of TBEV NS5 protein intensities quantified for each condition. D. Clustered heatmap of Pearson correlation comparing protein log_2_ intensity profiles for each pair of samples.

**Figure S3. Gene Ontology analysis of TBEV NS5 nuclear partners.** Categorical representation of Gene Ontology biological processes enriched among nuclear host factors interacting with the TBEV NS5 protein in infected cells.

**Figure S4. Further characterization of the interactions between NS5 and its partners using confocal microscopy and RNAse A treatment.** A. HMC3 cells were infected with TBEV at an MOI of 1. After 24 hours, the localization of selected TBEV NS5 partners was analyzed by immunofluorescence microscopy using specific antibodies (green). Infected cells were marked with antibodies against E or NS5 (red), based on species compatibilities. Nuclei were stained using DAPI (blue) and images were acquired using a confocal microscope. Acquisition settings were based on cells stained with secondary antibodies only (control). A. 293T cells were transfected for 24 h with plasmids encoding IGF2BP1-V5 and myc-IGF2BP3, or an empty vector as control (EV). Cell lysates were treated with RNase A or left untreated and were immunoprecipitated using V5 antibodies. Western blot analyses were performed on whole cell lysates (inputs) and V5-immunoprecipitates (IP) using the indicated antibodies.

