## Supplementary figures and images for "Nuclear activities and interactome of the NS5 protein of Tick-Borne Encephalitis Virus"

### Supplemental figures

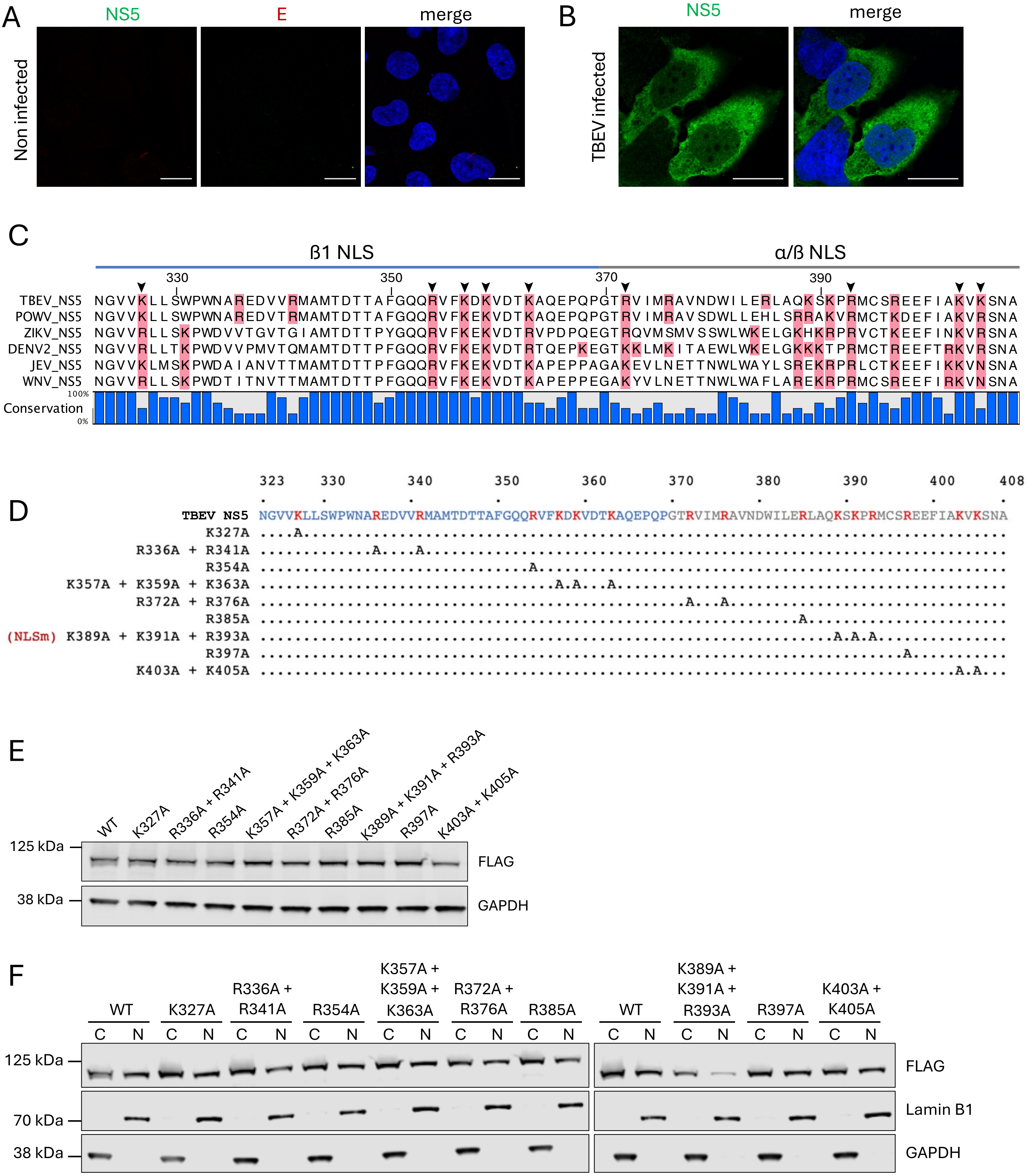

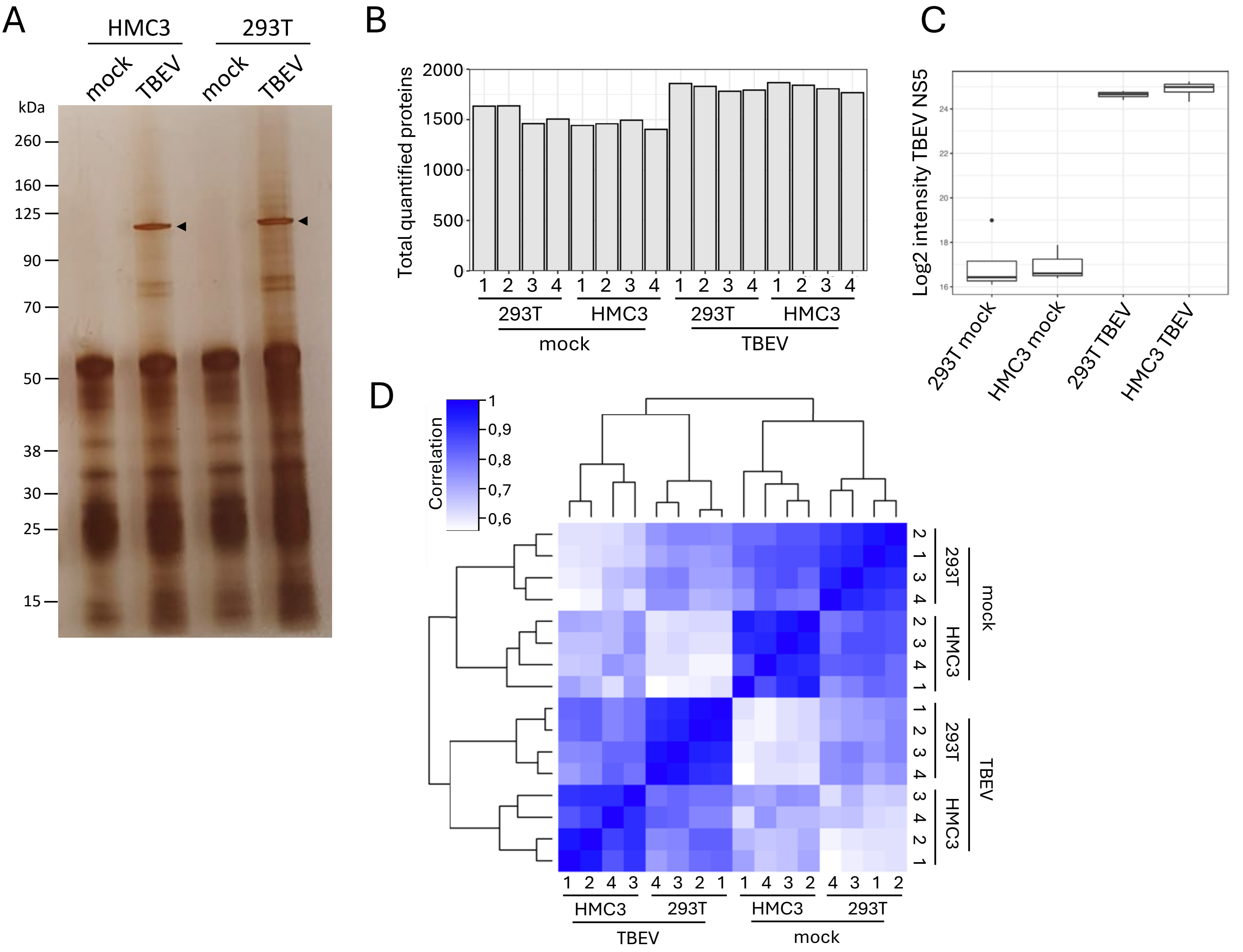

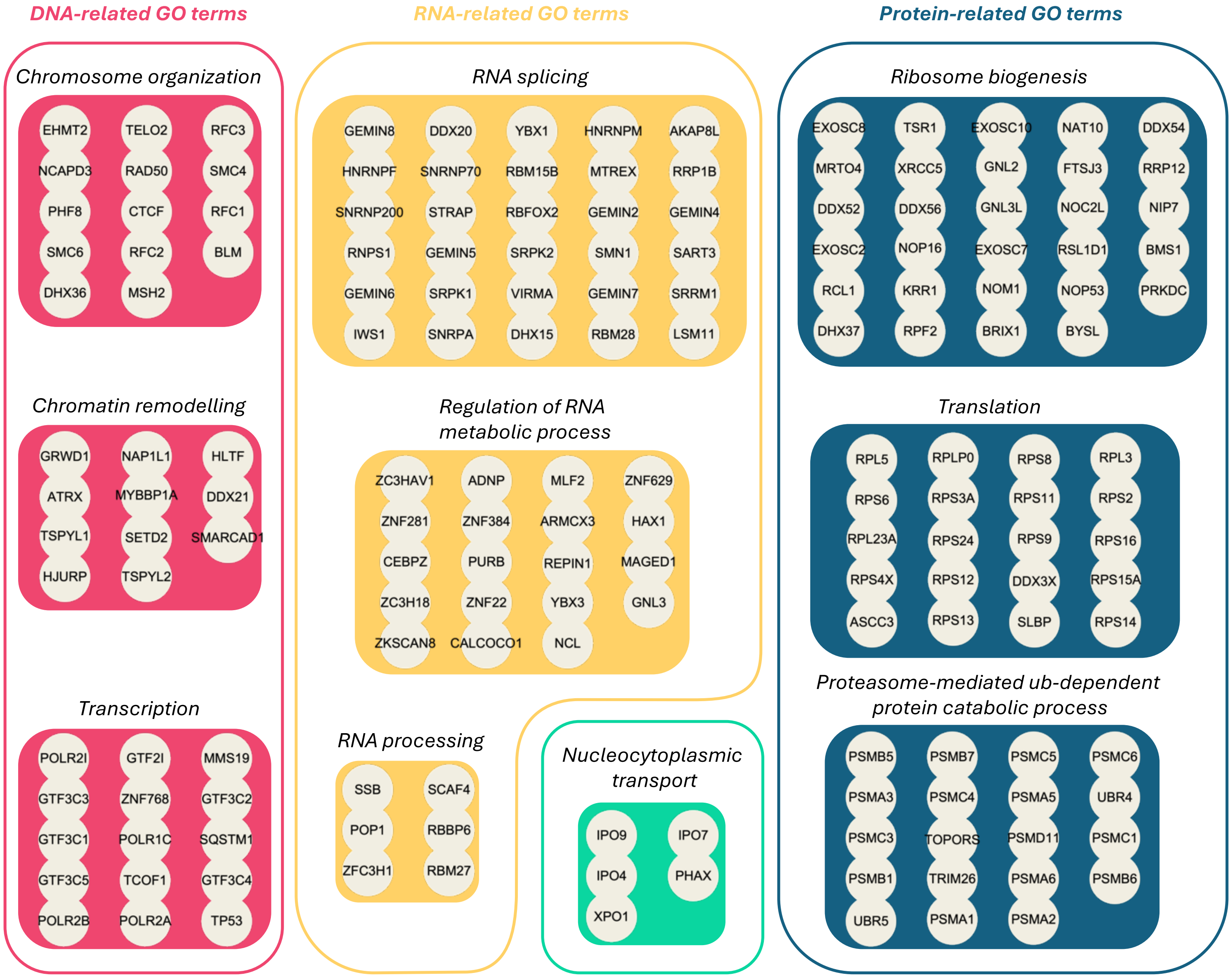

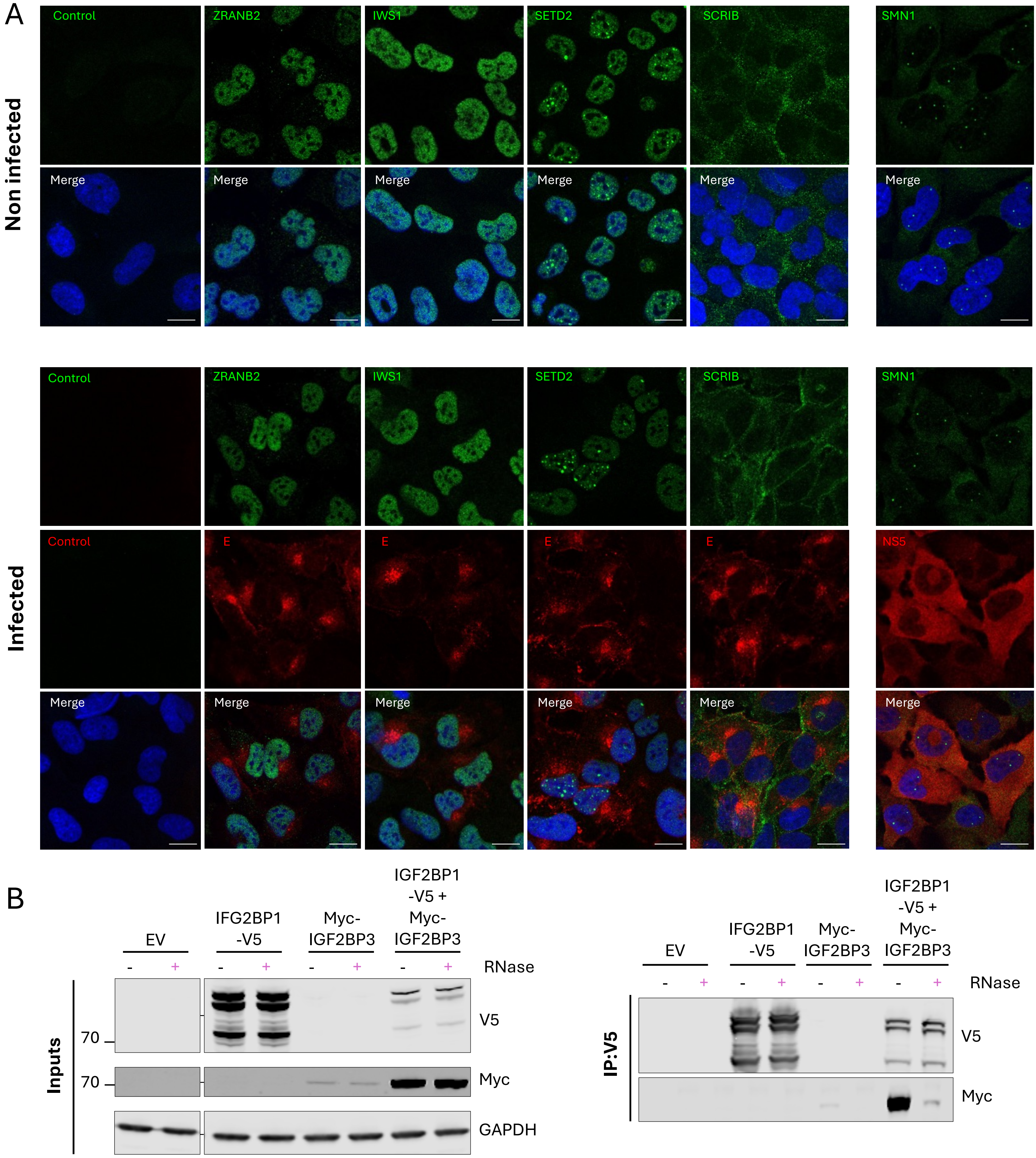
